# Fractal Features of Proto-Plant’s Origin and their Possible Consequences

**DOI:** 10.1101/083543

**Authors:** V.V. Galitskii

## Abstract

The extension of the sectional model of the spruce crown’s dynamics into diapason (0, 1) of the fractal parameter μ has demonstrated the existence of green biomass on branches of three orders in form of photosynthesizing (green) points. We investigated the growth of point sets on an interval as a model of the origin of proto-plants, which are formed due to endosymbiosis of cyanobacteria and protists. The fractal properties of the sets of evenly placed points and group sets were studied using the box-counting method. For the group sets, the character of dependence μ on the growing total number of points changes radically differently depending on whether the number of the points per group or the number of groups was fixed. As the host does not have the initial infrastructure needed for an increase in cyanobacteria per group, the first path is implemented and μ decreases from 1 to 0.25 when groups consist of two points per group. If and when the host develops necessary anatomical features (infrastructure), the second pathway is realized and μ grows to 1. The combined trajectory of μ initially demonstrates a slow growth of the size of the photosynthetic system and then an exponential growth after the development of the host’s infrastructure. Similar fractal peculiarity also characterizes trees and is an innate property of plants. Assumptions on the morphological recapitulation of proto-plant in higher plants’ ontogenesis (embryogenesis and seed germination) and also a possibility to fix the number of cyanobacteria per group are discussed.

## 1. INTRODUCTION

Morphological features are the first criteria that botanists pay attention to when studying plants. Although B. Mandelbrot has long noted the fractal nature of the architecture of trees and plants, this property is seldom taken into account by botanists when analyzing visual observation of plant architecture, more so in studies of the origin and evolution of plants. In this paper, an attempt is made to apply the basics of fractal geometry to study the endosymbiotic origin of plants, linking its green biomass *B* and size *H*

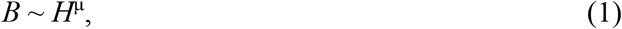

with help of the fractal parameter μ. These studies were inspired by an original concept about the sectional structure of trees (Galitskii 2006, Galitskii 2010; Barthélémy and Caraglio 2007) and a notion of virtual trees, thus allowing to implement this idea into a model of the vertical distribution of a tree’s green biomass. The original concept of the sectional structure of a tree and a model employs the known property of metameric (periodic) growth of a plant and the limitedness of its photosynthetic biomass (Galitskii 2010; Poletaev 1966).

In the sectional model of a tree, we used an image of coaxial virtual trees which are, at any particular moment, nested inside each other and periodically appear on the top of the real tree (Fig. 1). Each pair of such adjacent virtual trees forms a section of the trunk of a real tree. It is natural to assume that the dynamics of the *green* biomass of each virtual tree can be described by a model that is analogous to the real tree but with a parameter depending on the height of the appearance of the virtual tree (from the number of the corresponding section). The model of biomass dynamics of each section of the trunk is thus the difference of the green biomasses of the adjacent virtual trees forming the particular section (Galitskii 2006, 2010). In result, this model of a tree has demonstrated the acropetal denudation of a trunk of a freely growing tree observed in many species. Moreover, this model shows ultimate biomass distribution (with age) in the shape of an umbrella, being characteristic for palm trees and savannah trees (The Great Savanna … 2018), and also a number of biomass distributions along the height associated with species very distant from each other.

**Fig. 1.**
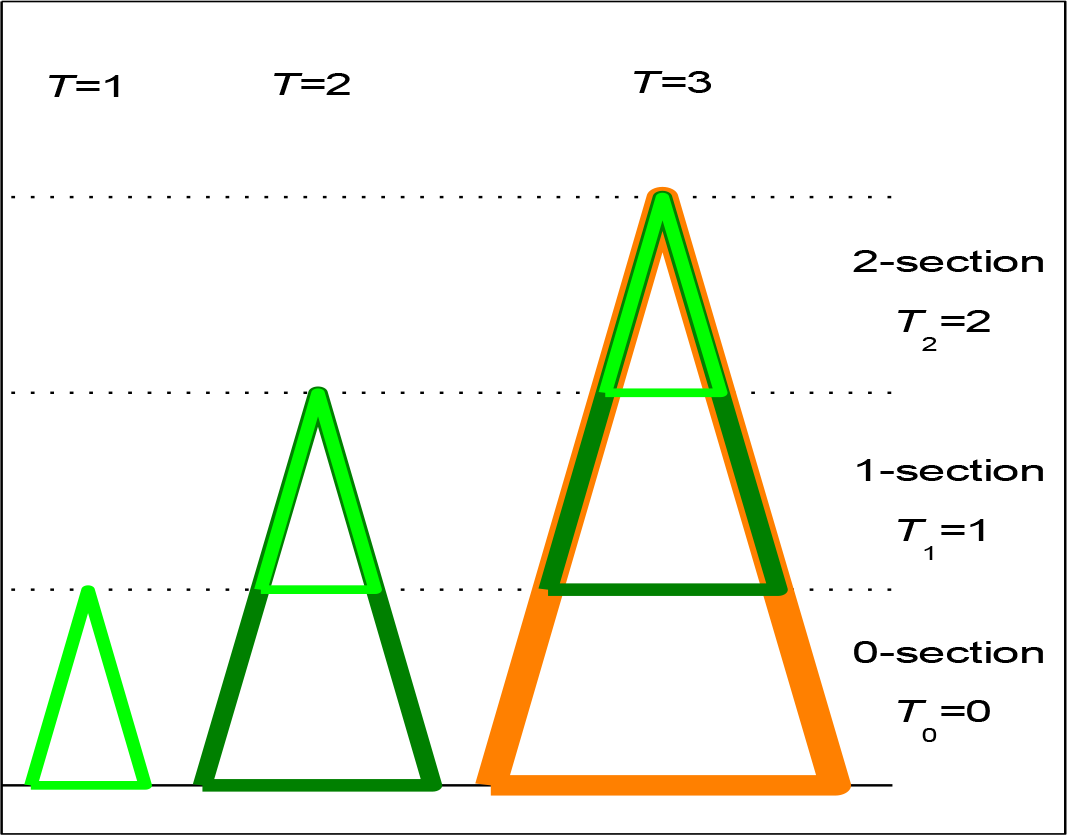
The scheme of the appearance and growth of sections and virtual trees. In the figure, at every age of a real tree, a triangle presents a virtual tree and a section is shown as a new triangle at the top of the real tree or as the difference of each pair of triangles of neighboring virtual trees.

Then we propagated the model to a system of *regular* branches of all orders that carry green biomass of respective sections of the tree (for example, spruce) in its ontogenesis (Galitskii 2013). The attempt of parameterization of this model of regular branches system of spruce using the data of lifespan (Tsel’niker 1994) of branches *t_D,j_* (*j* = 1, …, 4) of all orders (Table 1) has shown inadequacy of this model since such a model tree could have only branches of the first order, whereas the real spruce in the Moscow region has branches of four orders.

**Table 1.** Natural (Tsel’niker 1994) and modelled (Galitskii 2013) lifetime values *t_D,j_* for the *j*-branches of *Picea abies* (L.) Karst.

| Model variants | $t_{D,1}$, [year] | $t_{D,2}$ [year] | $t_{D,3}$ [year] | $t_{D,4}$ [year] |
| --- | --- | --- | --- | --- |
| **Natural data, sample 1** | **31** | **15** | **11** | **“7”** |
| Regular branches only | >350 | - | - | - |
| Regular branches + ID, #31 | >350 | 200.71 | 11.327 | 7.3063 |
| Regular branches + IV, #31 | 35.946 | - | - | - |
| Full model's values, #31 | 31.30 | 14.86 | 11.25 | 7.306 |
| **Natural data, sample 2** | **26** | **12** | **8** | **“5”** |
| Full model's values, #19 | 26.14 | 11.64 | 7.578 | 5.089 |

*Note:* Full model – the regular branches + initial deceleration (ID) + inter-verticillate branches (IV); natural data, samples 1 and 2, #19 and #31 – corresponding sets of values found with model parameters from Table 2; quotes – corrected natural data; an age of the branch is read off from the moment of appearance of appropriate tree section; tree section $i=10$ .

In previous publications (Kazimirov 1971; Treskin 1973; Kramer and Kozlowski 1979), two features of spruce growth were described, and in article (Galitskii 2013) two corresponding sub-models were added to the sectional model of the spruce branch system. The combined inclusion of their sub-models in the tree’s model resulted in a complete concordance with the real data. The individual inclusion of each sub-model had its own effect. The first sub-model (initial growth slowdown, Kazimirov 1971; Poletaev 1966) gave a complete qualitative correspondence to the real order of branching, but the quantitative discrepancy between the model and real lifespans of the branches all of the orders. Although the second submodel (inter-verticillate branches, Tsel’niker 1994; Treskin 1973; Kramer and Kozlowski 1979) did not provide individually realistic results, in combination with the first submodel it compensated the shortcomings of the first submodel. This distribution of the roles of the two mechanisms in the model allows us to assume that it corresponds to their positions in the evolution of the tree, and to think about the possibility of applying this model to the study of evolution.

In the work (Galitskii 2017) allometric parameter μ in Eq. (1) was interpreted as the fractal parameter. It allowed propagating the model of the tree’s branches system at all the real range of μ (0, 3). For modern spruce, parameter value obtained (Galitskii 2013) is μ≈1.8. It lies in the range [1, 2) that corresponds to a set of linear elements (conifer needles) according to notions of fractal geometry (Feder 1988). Fractal interpretation of expression (1) allowed to structure the analysis of the tree branch system model in accordance with the ranges μ [1, 3) and (0, 1) and apply the corresponding approaches for them.

The analysis of the model within the range μ [1, 3) shown interesting peculiarities of dependencies *t_D,j_*(μ). In Fig. 2 we show dependencies (trajectories) of lifespan *t_D,j_*(μ) of spruce branches of all *j*-orders in the real range of parameter μ for the combined model of a system of regular branches and initial deceleration of tree growth, i.e. when a proto-spruce did not yet acquire the mechanism of appearing inter-verticillate branches and can, therefore, be considered as an evolutionary precursor of modern spruce and, possibly, many coniferous trees (for more details, see Galitskii 2016, 2017).

**Fig. 2.**
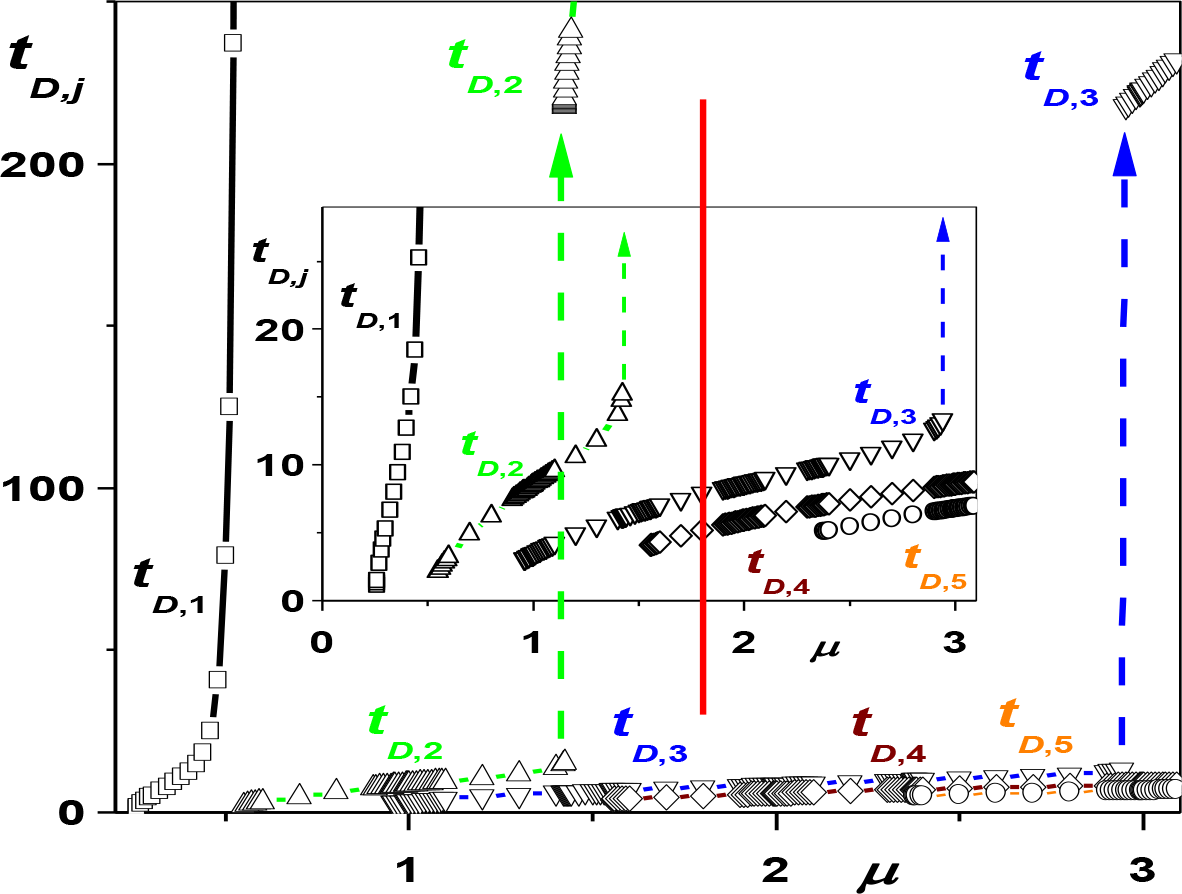
The model trajectories of the branch lifetimes of the orders *j*, (*j* = 1,…, 4) – *t_D,j_*(μ) [year]) in range μ (0, 3) for real spruce parameters (after Galitskii 2017). The inset shows an enlarged scale. The model (Galitskii 2010) of regular branches is supplemented with initial growth deceleration. Dashed lines indicate ruptures of trajectories; vertical red line indicates the united position (μ = 1.8) in both figures for the today spruce (Table 2).

**Table 2.** Parameters (Galitskii 2013) of full model of system of the spruce branches for the natural data (Table 1)

| Parameters | $A_1$, [year] | $r$ | $\mu$ | $\tau_w$, [year] | $w$ | $t_B$, [year] | $\beta$ |
| --- | --- | --- | --- | --- | --- | --- | --- |
| #31, sample 1 | 219.1 | 0.6649 | 1.830 | 8.994 | 2.637 | 8.700 | 1.235 |
| #19, sample 2 | 344.6 | 1.120 | 1.861 | 5.383 | 3.021 | 7.507 | 1.326 |

*Note:* $A_1$ – model parameter, $H_{\text{pl}}(T) = H_m \tanh(T/A_1)$ (Poletaev 1966); $r$ – parameter linking the maximum biomass $B_{m,i}$ of virtual $i$ -tree with height $H_i$ of its appearance, $B_{m,i} = B_m * f_0(x)$ , $f_0(x) = (1 - x)^r$ , $x = H_i/H_m$ ; $\mu$ – the fractal parameter in Eq. (1); $\tau_w$ and $w$ – model parameters of the initial growth inhibition $H(T) = H_{\text{pl}}(T) * (\tanh(T/\tau_w))^w$ ; $t_B$ and $\beta$ – model parameters of the inter-verticillate branches.

It is possible to note the presence of discontinuities of the first kind in dependencies *t_D,j_*(μ) for *j* = 2 and 3, and value μ ≈ 1.4 for the jump of 2-branch of spruce. As revealed in (Galitskii 2017), all ruptures disappeared when the combined model of the tree was supplemented with the sub-model of the mechanism of appearance of inter-verticillate branches (full model). Here we will not discuss the mechanism of the emergence of discontinuities and their role in the evolution of conifers (Galitskii 2016) and will focus on the range μ (0, 1). According to the ideas of fractal geometry, this diapason corresponds to sets of points (Feder 1988).

In this diapason, we have demonstrated in more details (Fig. 3), the dependencies of the lifetimes *t_D,j_*(μ) of spruce branches of the *j*-orders for sample #19 (Tables 1 and 2). It shows the features of the system of branches in the range of the parameter μ (0, 1) and corresponding to other parameters of spruce from Table 2 for regular model, supplemented by sub-model of the initial growth deceleration. In this range of μ, there are branches of three of the first orders, i.e., photosynthetic biomass is placed on a “tree” in the form of green “dots”, which is not observed visually for higher plants. In a study (Galitskii 2017) it was demonstrated that appearance of branches of the orders, *j* = 1, 2 and 3, in this range of μ is a consequence of the initial growth deceleration in the model.

**Fig. 3.**
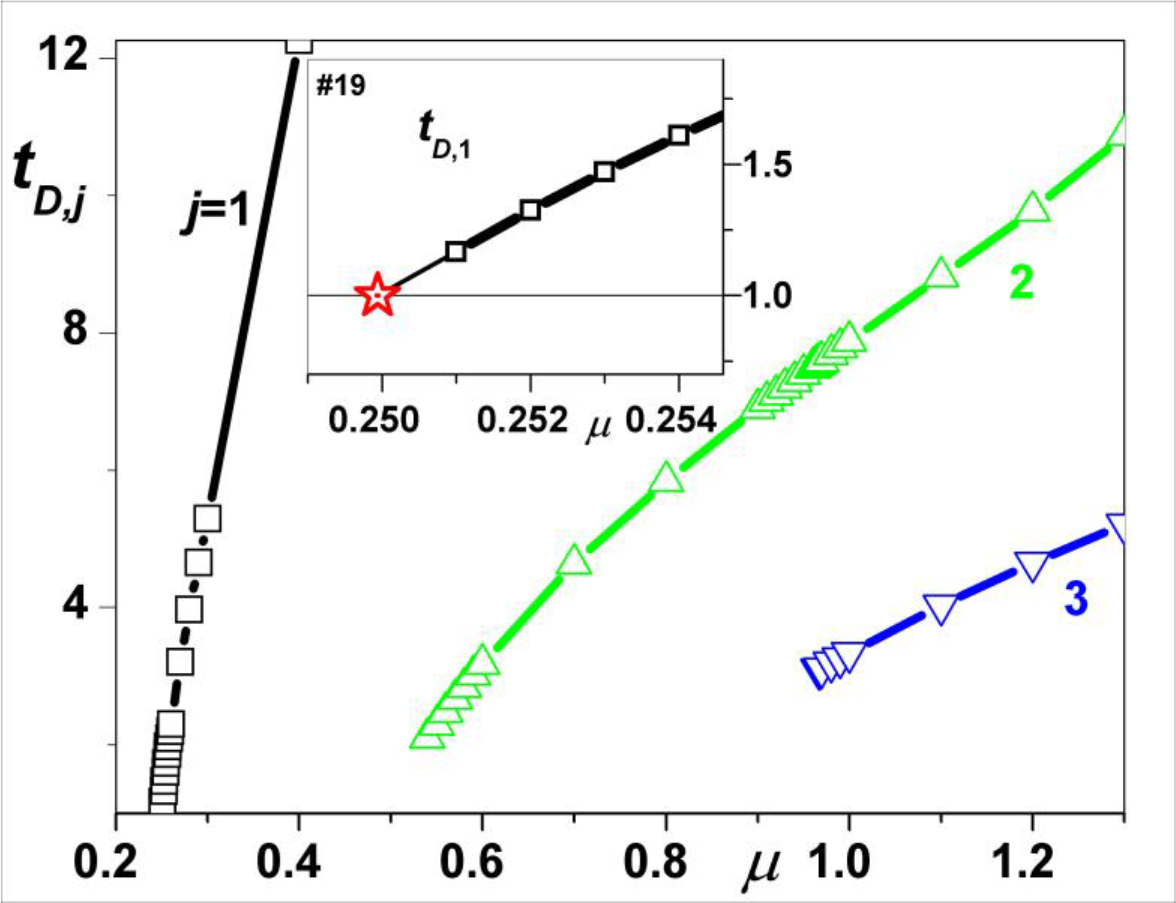
The dependence of lifetime of *j*-branches *t_D,j_*(μ) [year] in range μ (0, 1) ^11^. Other parameters are same as for real spruce. The model of regular branches is supplemented *only* with the submodel of initial growth deceleration (Galitskii 2013). The inset shows a 1-branch on a larger scale, asterisk – the appearance of 1-branch at μ = 0.24966 for the spruce parameter set #19, (Tables 1 and 2); (after Galitskii 2017).

## 2. A GROWING SET OF POINTS AND THE PROTO-PLANT’S ORIGIN

### 2.1. Photosynthetic system of plants as a set of points

The mechanisms of photosynthesis of higher plants and cyanobacteria are practically identical. This similarity is due to a history of the appearance of plants in the course of symbiosis and endosymbiosis of cyanobacteria and protists. According to the theory of Margulis (1981), chloroplasts of modern plants are descendants of cyanobacteria, which had undergone transformation after the organization of endosymbiosis with protists, in the course of evolution. Thus, “point” elements (chloroplasts), present in the cells of a green biomass of plants (Hanson and Kohler 2006), till now execute photosynthesis in plants. The features of placing a set of points, e.g., chloroplasts, in a space, affect the value of the fractal parameter and thus the efficiency of using sunlight in plant growth. We should note that a tree and a set of points, such as chloroplasts, correspond to natural objects of fractal geometry, which allows using its methods in the model directly (Feder 1988).

### 2.2. On properties of point sets

We assume that initial invasions of cyanobacteria into a cell of endosymbiosis host were sporadic and did not lead directly to compact placement, and that a protist is bigger than cyanobacteria by several orders. Therefore, we can estimate parameter μ for a set of *N_p_* points on an interval using the conventional algorithm of *box-counting* (Feder 1988). Placing points of a set, not in groups of points on interval but with uniform random placement or fixed steps, leads to μ ≈ 1.0, whereas placement in groups of points gives a value of μ <1. It should be noted that Cantor sets of points, which are often cited as a simple example from fractal geometry (Feder 1988), actually are group placements with μ <1.

For group sets of points at interval, it was shown that 1) the value of μ is practically independent of *A* – relative value of the off-duty factor for placement of groups; 2) μ depends on the quantity and the type of distribution of points in groups (Galitskii 2017).

We also found that growing group sets of points have a property which is important for understanding the regularities of the emergence and growth of plants from the proto-plant to the modern giant sequoia. This property is as follows: The growth of the total number *N_p_* of points in the group set (*N_p_* = *N_g_* * *n_g_*) in the simplest case can be organized in two ways: 1) an increase in the number of groups *N_g_* for a fixed number of points *n_g_* per group and, 2) an increase in the number of points per group for a fixed number of groups. The behavior of the fractal parameter μ with increasing *N_p_* for these two variants is radically different. We will consider these options in the case of the appearance of the proto-plant below.

## 3. RESULTS

### 3.1. Trajectories μ(*N_p_*=*N_g_*\**n_g_*) for fixed *N_g_* and fixed *n_g_* are different

In Fig. 4 we demonstrate examples of dependences of μ(*N_p_*) (small empty quadrates, solid lines) for some fixed values of *n_g_*=*n*_*g*,fix_. The parameter μ decreases from 1 to ≈ 0.25 with the growing number (beginning from 2) of groups *N_g_* with a minimum fixed number *n*_*g*,fix_ = 2 of points per group. The same figure shows examples of dependencies of μ(*N_p_*) with increasing *n_g_* (beginning from 2) for several fixed values *N_g,fix_* (the same small quadrates, dashed lines). There is a weak increase in the parameter μ with the number of points in group *n_g_* for a small fixed number of groups *N_g_* (≥ 2). When the value of *N_g,fix_* is large, the parameter μ initially increases rapidly with *n_g_* and then slows down; remaining at a value of slightly less than 1. The abscissa in Fig. 4 shows the evolutionary “ersatz-time” *T*_2_ as the total number of doublings of the corresponding active (not fixed) variable – *N_g_* or *n_g_* (chloroplasts reproduce by dividing in half). The value of *T*_2_ is the number of doublings of cyanobacterial symbionts *N_p_,* it demonstrates the state and direction of the process and is not an evolutionary time, which is much larger and controlled by the rate at which the infrastructure is created by the host.

**Fig. 4.**
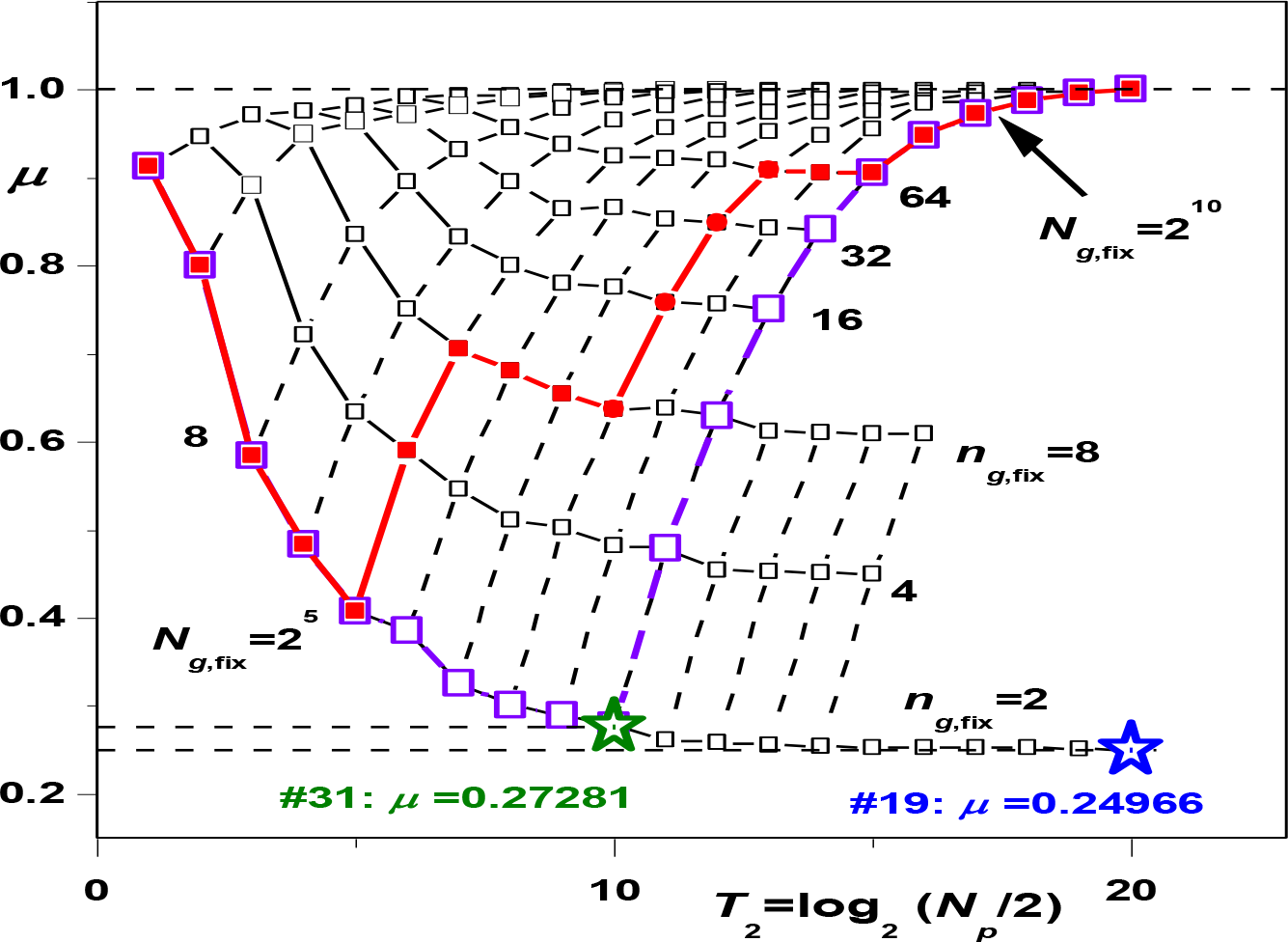
Two-dimensional images of three-dimensional trajectories of dependencies μ(*N_g_*,*n_g_*) for group placings. Two types of trajectories, viz., waning trajectories of μ for a fixed values *n*_*g*,fix_ (thin solid lines, small empty quadrates) and growing trajectories for a fixed values *N*_*g*,fix_ (thin dashed lines), which accordingly pass through the same points (empty small quadrates) with ordinates μ(*N_g_*,*n_g_*) and integer abscissas log_2_(*N_g_*\**n_g_*/2). Two examples of proto-plant evolution are made from elements of two mentioned types trajectories (by thick solid lines with filled small quadrates and thick lines with empty big quadrates): both begin at different places of trajectory *n*_*g*,fix_ =2 and go on different paths to μ=1. First thick trajectory demonstrates a possibility of regressive evolution (decreasing μ) until the host cannot create the necessary infrastructure to go to the next growth trajectory. The off-duty factor *A* = 0.005 shows the ratio of the length of a group to the size of step for the placement of the groups at the interval with a step. The empty green and blue asterisks — μ#19 and μ#31 (see an asterisk in the inset in Fig. 3 for #19 as an example how to find these values μ) correspond to natural spruce data (Tsel’niker 1994) and their modeled interpretations (Galitskii 2013) (Tables 1 and 2). *T*_2_=log_2_(*N_p_*/2), *N_p_*=*N_g_*\**n*_*g*,fix_ or *N*_*g*,fix_\**n_g_* (see text) is evolutionary “time” measured by the number of doublings of *N_p_* along a trajectory. *T*_2_ is not real time of evolution.

### 3.2. Fractal trajectories of proto-plant’s evolution

Endosymbiosis, when cyanobacteria exist within the protist, begins with a symbiosis of the protist and bacteria that are located in an environment outside the protist. Symbiosis is a mutually beneficial cohabitation, in which there is primarily a material interchange. Cyanobacterium, which got into the protist, should consume and excrete and periodically breed, and the host being benefited in the symbiotic production has to have the necessary morphological adaptations to create infrastructure linking the symbionts and the host in a united system. Initially, these symbiotic relationships are either absent (primary endosymbiosis) or insufficient to maintain more than a pair of symbionts in the group (endosymbiosis after symbiosis). In the first case, the invasion of cyanobacterium results in its death or digestion, in the second case a gradual transition from symbiosis to limited endosymbiosis is possible. Therefore, the evolution of such an endosymbiosis in the host with the limited infrastructural capabilities can begin to go only along the extensive path of randomly increasing the number of such local groups with a minimal number of cyanobacteria per group. This is expressed by an increase in the photosynthetic products but as it is seen in Fig. 4, with a decrease in μ together with the efficiency of making use of the available light energy. So, when cyanobacteria have an opportunity of restricted dividing (*n_g_* ≤ 2), and restriction (if any) for the number of groups *N_g_* is removed, evolution begins along trajectory with *n*_*g*,fix_ = 2 (Fig. 4) tending to μ≈0.25.

In the course of the natural evolution of both organisms involved in the endosymbiosis, morphological and other adaptations appear which cause an increase in the number of photosynthetic units in the group and of total photosynthetic production. Such changes inside cyanobacteria, as shown by micromorphological studies (Ting et al. 2007), may have begun even before cyanobacteria were included in the endosymbiotic trajectory of evolution. As a result, the system of endosymbiosis was able to move in a certain growth path μ. If the *n_g_* restriction is retained, then it again leads, as can be seen from Fig. 4, to another waning trajectory μ with some larger *n_g,fix_*. Such stages of evolution could occur more than once.

In the course of evolution, when at some “time” *T*_2_, immediately or gradually, the restriction of cyanobacterial division inside groups is removed, the evolution of proto-plant moves to appropriate trajectory *N*_*g*,fix_ (Fig. 4), which leads to an increase in the magnitude of μ to the value of 1. When fractal parameter μ reaches to 1, the situation already corresponds to the sets of linear elements, i.e., piece-wise linear or continuous filling of the interval (considering the real size of green “point”). In Fig. 4, we show this procedure in the assembly of several combinations of fixed values *n*_*g*,fix_, and *N*_*g*,fix_.

Thus, during the evolution of proto-plant, dependence *μ*(*T*_2_) of the fractal parameter must proceed from 1, reach to a minimum value and again increase to a value of 1. In this scenario, of course, more profound detailing is possible, depending on the symbiotic “experience” of protist and cyanobacteria. For example, a trajectory could start with *n*_*g*,fix_= 4 or go to the growing branch, depending on the value of *N_g_* at which the trajectory changes its type, as shown in Fig. 4.

Such types of changes can occur repeatedly, but the general character of the trajectory of the proto-plant does not alter. The countability of sets of variants of these evolutionary trajectories may be noted. Each of these variants move proto-plants into a state with μ = 1 in different evolutionary times, and this could consequently lead to different histories of plant evolution. In Fig. 4, two positions at the trajectory *n*_*g*,fix_ = 2 corresponding to values *μ* (#19 and #31, Table 1), obtained for two sets of natural values^1^ of *t_D,j_* for spruce (Tsel’niker 1994), are marked by asterisks. It is evident that the proto-spruce (#19) spent approximately twice as much “time” on evolution than the time spent by the proto-spruce (#31), and this was reflected in their “descendants” (see Galitskii 2017: Fig. 9).

### 3.3. Initial growth deceleration of a proto-plant

As seen from previous data, the green biomass (i.e., the number of cyanobacteria involved in endosymbiosis) is the leading variable in the evolution of a proto-plant. Therefore, “turning over” the ratio Eq. (1) and using the obtained dependence μ(*T*_2_) (Fig. 4), we can calculate the “dynamics” of the characteristic size *H* of the system of points–*H* (*T*_2_) ∼ (*N_g_* (*T*_2_)\**n*_g_ (*T*_2_))^1/*μ*^(*T*2). In Fig. 5, we demonstrate examples of dependences of *H* (*T*_2_) of weak growth at initial *n*_*g*,fix_ stage followed by the exponential rise at the *N_g,fix_* stage.

**Fig. 5.**
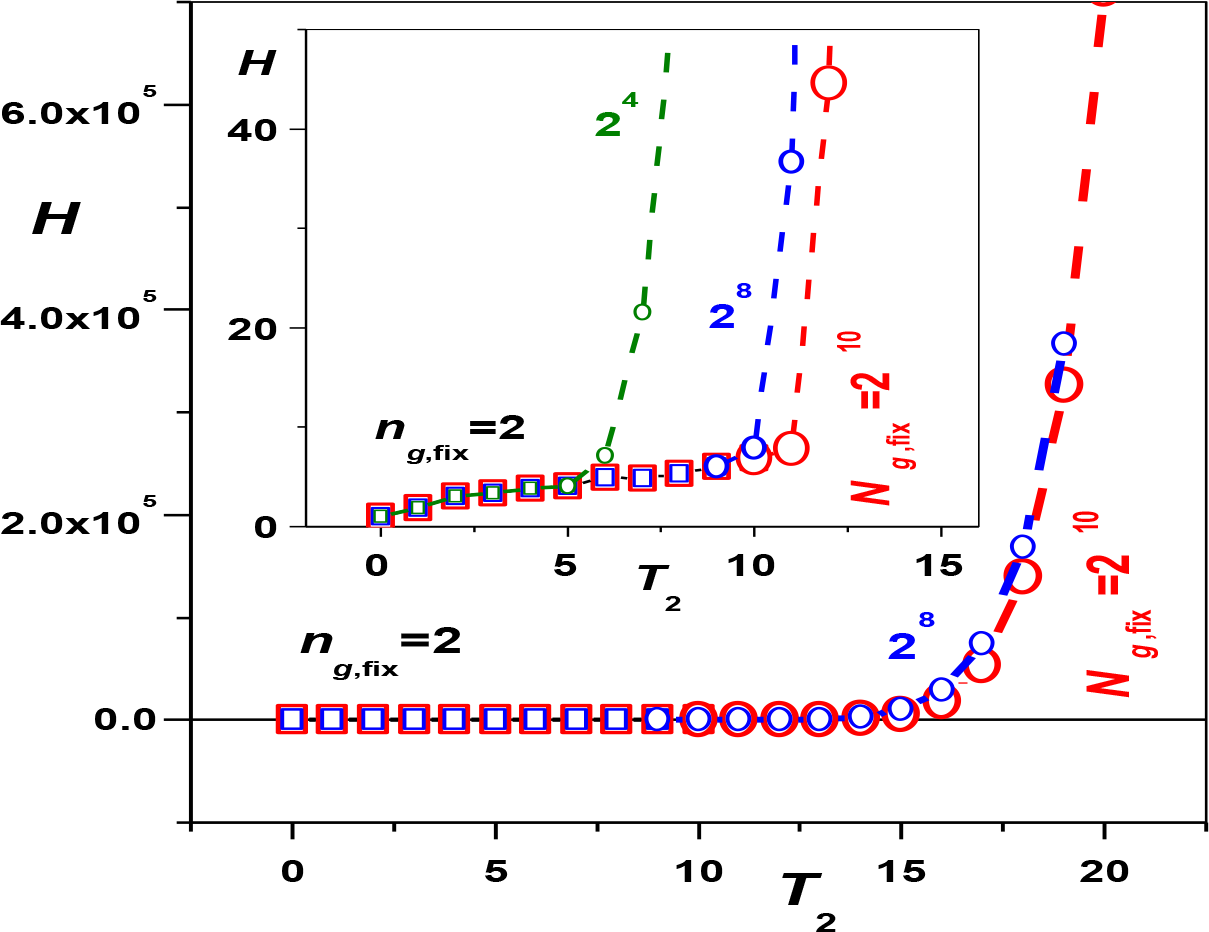
The initial deceleration of the proto-plant’s size growth during the endosymbiosis. The dynamics of size *H*∼*B*^1/μ^ of the set of green “points” with increasing their total number *N_p_*=*N_g_*\**n_g_* (and biomass) by way of doubling the number *N_g_* of groups or “points” *n_g_* per group (biomass and size units are the mass and size of single cyanobacterium); the combined dependence of fractal parameter μ(*T*_2_) used (see Fig. 4). The inset shows an enlarged scale: weak growth in the first part during movement along trajectories μ(*n_g,fix_*,*N_g_*) at fixed the number points *n_g,fix_* per group with the increase in the number of groups of *N_g_* (squares). In the second part at fixed number of groups *N*_*g*,fix_, during movement along trajectories μ(*n_g_*,*N*_*g*,fix_) with the increase in the number of *n_g_* points per group (circles, dashed) – exponential growth *H*. *T*_2_ – see Fig. 4.

## 4. DISCUSSION

### 4.1. Are there recapitulations of the proto-plant in the ontogeny of modern plants?

According to modern evolutionary theory (biogenetic law), sometimes, in embryonic development, the stages of evolutionary history can be reproduced, i.e., one may see so-called recapitulations. Usually, the obvious visual similarity of a certain stage of embryogenesis of a contemporary organism and ontogenesis of its evolutionary ancestor is considered to be a recapitulation.

In this sense, for example, such “point recapitulations” that visually and sufficient “strictly” repeat the model situation of proto-plant’s emergence (as at the beginning of the range μ <1.0, in Fig. 3) may apparently be detected in modern plants twice.

At the first time in dicotyledons, in the initial stages of embryogenesis, an embryo of a seed begins to turn green due to the transformation of the existing colorless etioplasts (leucoplasts) into chloroplasts (Zhukova 1992), in the course of transition from globular to the heart phase of embryogenesis (Fig. 6). The form (two cotyledons) of the embryo in the heart phase (Serebryakova et al. 2006) and placement of chlorophyll (Tejos et al. 2006) at initial phase may be associated with the appearance of “branches” of the first order of proto-spruce at μ≈0.25 (Fig. 3). Obviously, there cannot be a complete coincidence. Proto-plant has appeared and propagated in the absence of the appropriate infrastructure and corresponding elements of the genome in the host, whereas the seeds of a higher plant in early embryogenesis already have all this in some form or other.

**Fig. 6.**
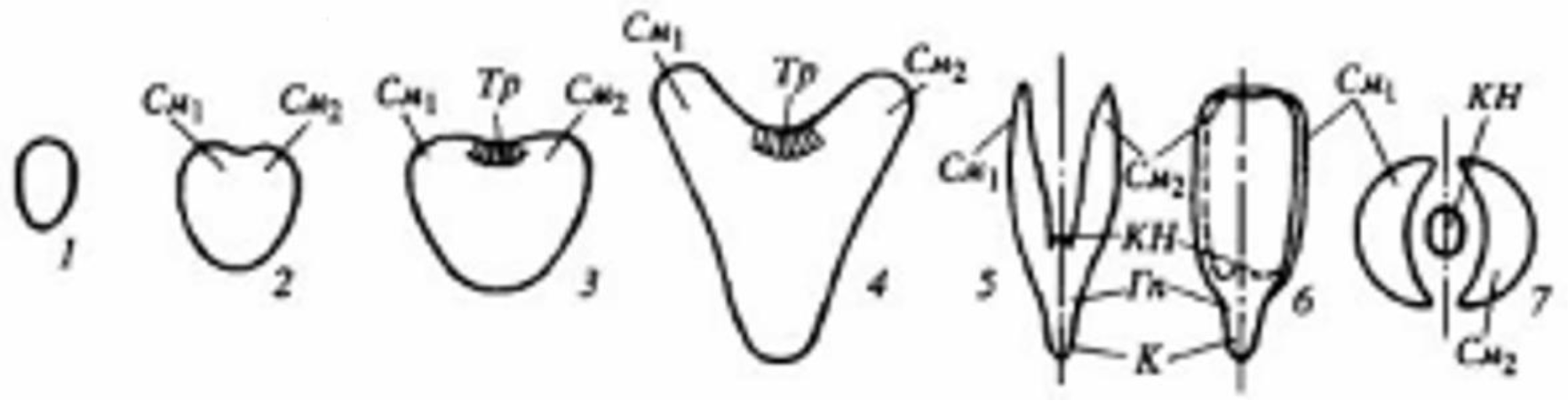
The scheme of the development of dicotyledon embryos (after Serebryakova *et al.* 2006: 144, Fig. 70). 1-4 – early development of the embryo (1 – globular phase, 2 – 3 – heart phase, 4 – torpedo phase); 5, 6 – fully formed embryo; view of two mutually perpendicular planes; 7 – cross-sections in plumule’s embryo. *C_m1_* and *C_m2_* – cotyledons, *T_p_* – growth point (before forming the cone of growth *KH*); *Γn* – hypocotyl; *K* – root. The dashed line shows plane of symmetry. The heart and torpedo phases demonstrate the appearance of cotyledons, which are analogs of the “branches” of the first order, appearing in places marked with asterisks (Fig. 4).

According to the model, we can assume that embryogenesis moves along a composite trajectory (of course, far more rapidly than the proto-plant appeared and evolved), which is essentially similar to that shown in Fig. 4, taking into account the differences in their starting positions and structures. The placement of fixed number *n_g_*,_fix_ of plastids per group (Fig. 4, empty squares) during the growth of total number, in the globular phase of embryogenesis (Fig. 6, 1) before and after the transition to heart-shaped phase (*2*, *3*), possibly could be a distinctive feature of recapitulation (if any). The result of plant embryogenesis is the appearance of typically yellow, mature dry seeds, which have no active chlorophyll, as chloroplasts turn into colorless leucoplasts (plastids) during the maturation of seeds (Zhukova 1992), and thus the seeds are able to wait for appropriate conditions for germination.

At the second time, apparently, recapitulation occurs during germination of mature, dry, yellow seed of a plant, when the embryo with developed cotyledons begins to turn green as a result of the transformation of existing colorless leucoplasts into green chloroplasts. Typically, this natural process of increasing the number of active chloroplasts expresses in the normal ontogeny of a plant. However, for some economically important crops, ripe seeds sometimes remain green. It is unclear why such seeds, having additional (extra) chlorophyll, show poor germination and give low yields and quality of products of oilseeds (Nakajima et al. 2012).

If we assume that the situation at the beginning of germination of a seed and activation of “systemic” chlorophyll corresponds to the proto-plant’s model, then the role of “non-systemic” or excess chlorophyll becomes clearer. It is evident that the activation of chloroplast in an embryo, having appropriate infrastructure, is controlled by the genome. “Non-systemic” chloroplasts are not consistent with the infrastructure of an embryo, are located quite randomly and generate extra “non-systemic” oxygen that can damage cell structures (Nakajima et al. 2012). Such chloroplasts may adversely affect the process of germination and further ontogeny. Therefore, not only quantity but also the spatial placement of chloroplasts may be important for germination process to occur.

Thus, in the ontogeny of seed plants, two situations of recreating photosynthetic system could formally resemble, at least morphologically, a model situation of the emergence of a proto-plant. Both seeds and proto-plants, use the same elements that are in one of the two states – leucoplasts and chloroplasts and presumably use similar mechanisms to switch from one state to another.

Apparently, in connection with the present model, it would be advisable to conduct experimental studies using fluorescence confocal microscopy (Tejos et al. 2010) together with the model analysis of placement and the transformation of leucoplasts and chloroplasts in embryos and sprouts of wild and related mutants of Arabidopsis in early embryogenesis and in germination.

### 4.2. About initial deceleration the growth of modern plants

From the foregoing, it follows that an initial growth inhibition exists at the stage of formation of a proto-plant, i.e., it is the foundation of evolution leading to higher plants. Poletaev has shown (1966) that the model of isometric growth of a tree (i.e., without an initial deceleration), parameterized using natural data on the growth of oak (Molchanov 1964) describes sufficiently precisely the height growth of the mature tree (Fig. 7*a*). However, this model also showed a substantial difference in the initial growth – real growth begins noticeably later than the model indicates. In Fig. 7*b*, we demonstrate a slight modification of a model tree growth’s dependence, which was satisfactorily applied to a similar problem of fitting model values to natural data of the lifetime of spruce branches of all orders (Galitskii 2013, 2016) for all the ages. The quality of approximation may be seen when the natural and modeled values *t_D,j_* in Table 1 and values *A_1_* and *τ_w_* in Table 2 are compared. It is interesting to note that such slight fractal-induced modification has given branching of the high orders to trees (Galitskii 2013), and for forests and plants in general, beauty and charm.

**Fig. 7.**
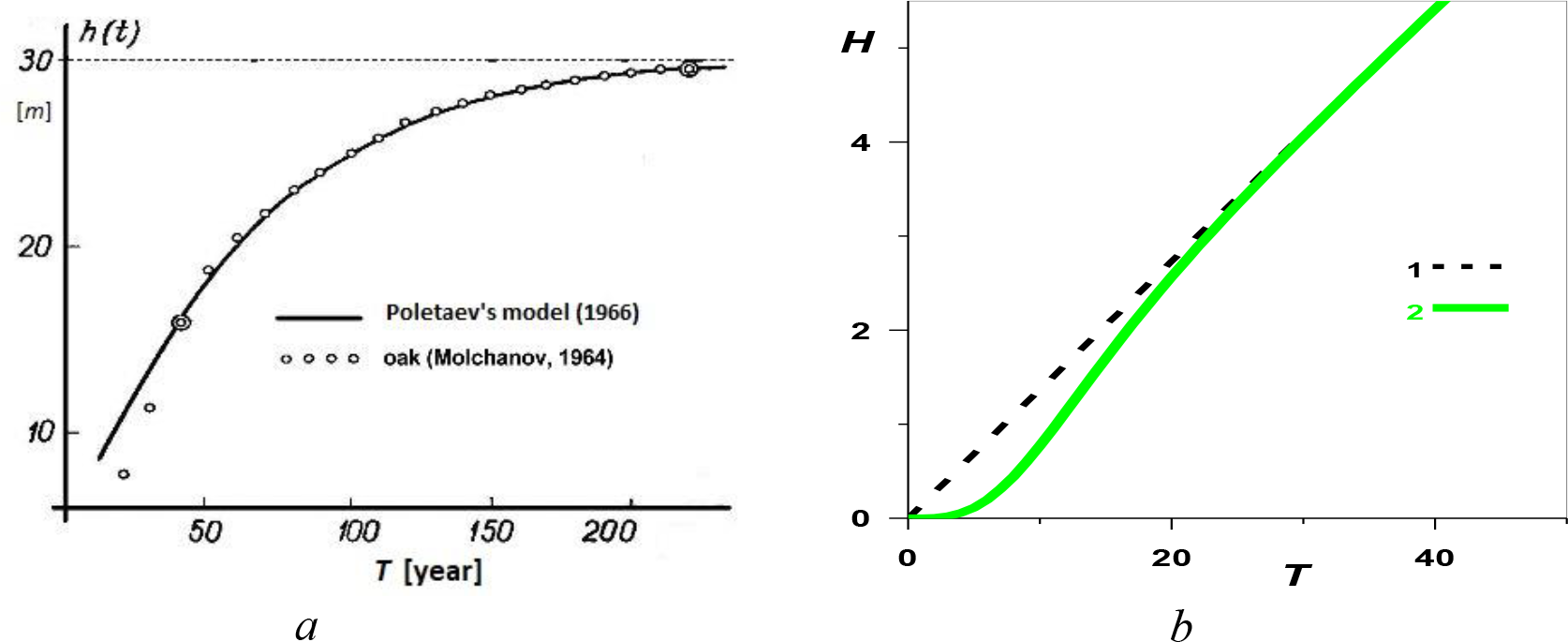
The initial deceleration in reality and its model interpretation. (a) Model dynamics of tree’s isometric growth, solid line – Poletaev’s model (1966) (see note to Table 2) and natural data (Molchanov 1964) (circles) for the oak tree, double circles – data used for parameterization of Poletaev’s model. Very good agreement of (∼tanh(*T*/*A_1_*)) in mature age, and the substantial differences in young age. (*b*) Modification (solid green line) of Poletaev’s model (dashed) with the multiplier (tanh(*T*/τ_*w*_)^*w*^ which is sufficient for the proper agreement for branches lifetimes (see Table 1).

The example in Fig. 8 demonstrates an initial deceleration followed by the acceleration of height growth and gradual increase of the maximum order of the spruce branches (Serebryakova et al. 2006) for young specimens of several first ages. The figure shows the spruce age and by numbers —the age [year]—when the relevant section of the stem appeared. This picture is consistent with previous data (as depicted in Fig. 3), and with the results of an earlier work (Galitskii 2013). In the first four years, there are no branches and biomass (needles) located on the stem. In the fifth year, there are branches of the first order (at μ ≥ ≈0.25, Fig. 3). The branches of the second order appear in the sixth year (at μ ≥ ≈0.55). In the seventh year, the branches of the third (at μ ≥ ≈0.9) and fourth order (at μ≥ ≈1.5, Fig. 2) and inter-verticillate branches of stem section (at μ ≥ ≈1.4) appear.

**Fig. 8.**
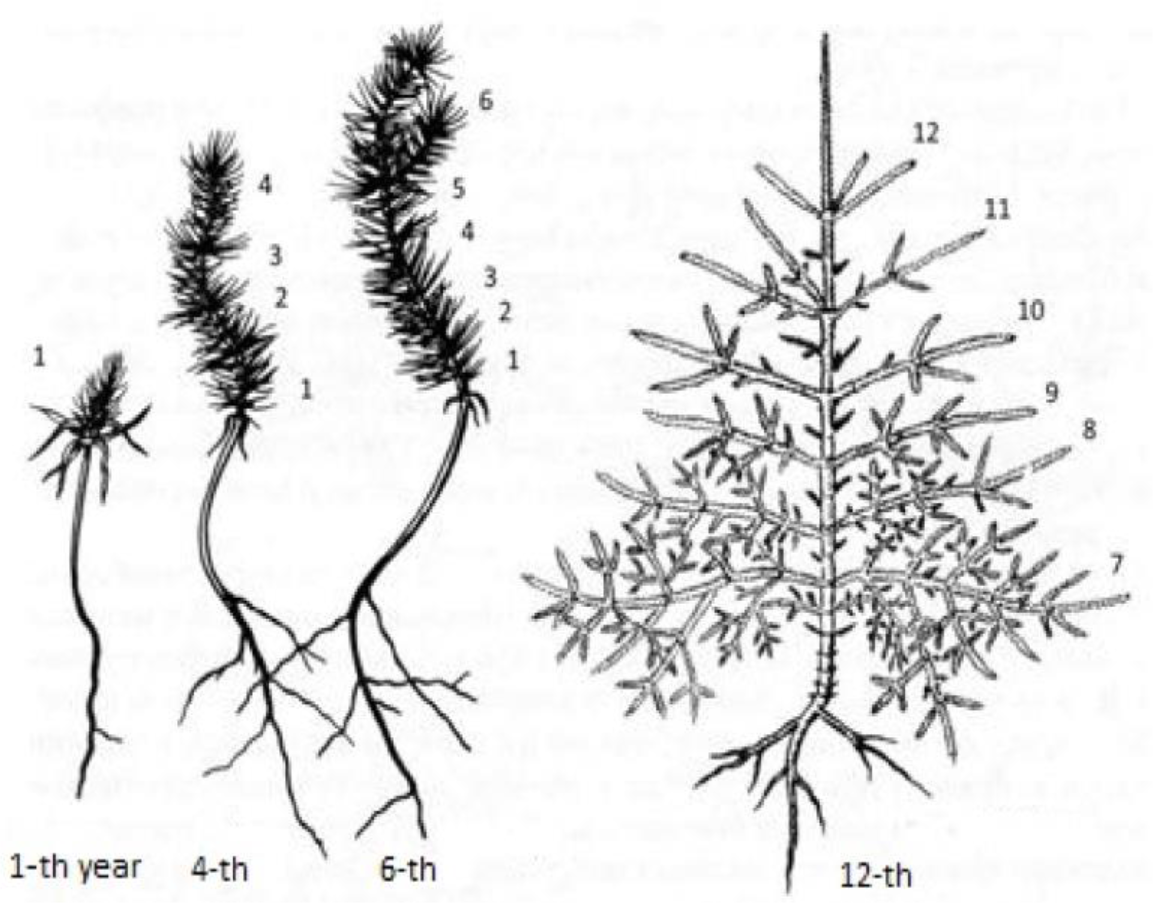
Formation of the branch system of spruce of young age, μ ≥ 1, see text (Serebryakova et al. 2006: 298, Fig. 186).

It is evident that there is no visual correspondence between juvenile stage of spruce and the emergence of the proto-plant. However, both show qualitatively similar dynamics of characteristic sizes of placing their photo-synthesizing biomass in space. The initial growth retardation after the exposure of a plant to light is similar to that for proto-plant. In agreement with Fig. 3 (and a previous study Galitskii 2016), this deceleration coordinates the existing material capabilities of a plant (i.e., its green biomass) and placement of its biomass in space, described by structural parameter μ, linked in this case to the geometry of the system of regular branches.

We may note that in the monograph (Kozo-Polyansky 1937), the author, using extensive literature data, discussed the features of the biogenetic law in botany, and also numerous testimonies of manifestations of recapitulations, which are possible not only for the morphological traits of organisms but also for paleontological, biochemical, and their other traits.

### 4.3. Plausible reasoning

Interpretation of growing group sets of points in an interval (Sec. 3.2) as a model of the endosymbiotic origin of a proto-plant raises questions that may not have direct answers. However, proceeding from the specifics of the topic and relying on the idea expressed and discussed in Polya’s monograph (1954) “Mathematics and plausible reasoning”, we may at least try to formulate some questions.

Firstly, the most pertinent question is how did the first cyanobacterium appear in a host’s body? One of the hypothetical ways described in the literature (for example, Martin 2001) is that the heterotrophic host envelopes a prey with its body, forming a capsule within itself. For the encapsulated organism, the capsule had the same order of boundary layers of the host as it had in a preceding symbiosis. In other words, the capsule contained the cyanobacterium is surrounded by the same symbiotic environment, which it had previously encountered. The position of internal compartments of the host (in particular, contractile vacuole and mitochondria) also does not change topologically, relative to the captured cyanobacterium. The capsule is arranged in a manner similar to a digestive vacuole of an amoeba and, generally speaking, could give rise to a future chloroplast. Some species of the proto-amoebae could serve as a host for endosymbiosis (Mereschkowsky 1905; Hausmann and Hülsmann 2003; Kutschera. and Niklas 2005). Comparative genetic analysis (Nozaki 2005) revealed the presence of a common ancestor in Amoebozoa and Green Plants, and that the plastids of the latter were obtained as a result of primary endosymbiosis.

In continuation of this question: how did the bacterium, which found itself in the digestive vacuole, avoid being digested? We can assume that the main products of cyanobacterial metabolism, viz., carbohydrates and oxygen, could have played a role here. It can be speculated that during previous symbiosis relations, the digestive enzymes of the host had been “adapted” to the cyanobacterial products.

Secondly, the described fractal nature of the appearance of endosymbiosis and the beginning of the evolution of the proto-plant require a group placement of cyanobacteria. Moreover, as the model shows, during very long intervals of host’s evolution, restricting the number of bacteria per group must go together with the increase in the number of groups of cyanobacteria (and the total number of bacteria accordingly). Cyanobacteria propagate by dividing in half (modern bacteria divide in < 24 h). Therefore, after reaching a certain number, one bacterium of each daughter pair must die off. The question of how this occurs is beyond the scope of the article (see as an example England 2013). However, we can make a note of a known example of modern endosymbiosis, with infusorium as a host, where the number of chlorella-symbionts is variable and restricted (Margulis 1981; Eskov 2008).

To get answers to these and many other questions, careful observations and, if possible, experiments with prospective candidates for participation in such endosymbiosis (Wöstemeyer et al. 2016) would be useful. Given the properties of the biological material, it is almost impossible to find remnants of participants and the sceneries of these ancient evolutionary occurrences. Probably the only way to analyze such phenomena is to go through a verified path of plausible reasoning and analogies: “Anything new that we learn about the world involves plausible reasoning, which is the only kind of reasoning, for which we care in everyday affairs. … You have to combine observations and follow analogies; you have to try and try again.” (Pólya 1954).

## 5. CONCLUSION

The expansion of the dynamics model of the green biomass of the spruce branch system (Galitskii 2013) into its fractal “past” (Galitskii 2017) – the “point” range of the fractal parameter μ (0, 1) showed the existence of the first three orders in the proto-spruce’s branches, i.e. presence of a certain geometric structure of the placement of green biomass in the form of point sets. The placement’s photosynthetic efficiency of this biomass in space is described by the parameter μ. Based on the theory symbiogenesis by Margulis (1981), the properties of group point sets was investigated as a model for the emergence and evolution of a proto-plant. It was found that growing group point sets, characterized by the number of groups *N_g_* and the number of points in the group *n_g_* and the total number of points *N_p_* = *N_g_* * *n_g_,* demonstrate a radically different behavior of μ with increasing *N_p_,* depending on whether *n_g_* or *N_g_* is fixed. Being applied to the formation of endosymbiosis of a host (protist) and cyanobacteria this property is naturally to associate with the initial absence or presence of the host infrastructure necessary for the functioning of the system which uses the material exchange between the symbionts. Initially, the host does not have the necessary infrastructure and therefore the minimum *n*_*g*,fix_ = 2 is fixed, *N_g_* grows and μ decreases. During the development of endosymbiosis (evolution of the proto-plant), when the infrastructure is created, *N_g_* is fixed accordingly, *n_g_* and μ grow. Thus, as *N_p_* increases for *n*_*g*,fix_ = 2, the trajectory μ goes from μ = 1 to a minimum of μ≈0.25 and then again increases to μ = 1. There may be more than one such minimum. It should be noted that the “moments” (μ values) of the appearance of first-order branches (asterisks in Figures 3 and 4), obtained by the method shown in Fig. 3, get on the same trajectory, *n*_*g*,fix_ = 2, of initial decrease μ (Fig. 4), calculated by a method in no way connected with the first method, which indicates the essential (informal) bond of the two models – of spruce and the appearance of the proto-plant.

The presence of intervals with a decrease in μ is expressed in inhibition of the growth of the size of the photosynthetic system. The very beginning of the evolution of the proto-plant is characterized by a slow growth in size, which is replaced by an exponential growth after the transition to the growth trajectory μ. This initial slowdown in growth and development is also characteristic of modern plants, which is also associated with the fractal spatial distribution of photosynthetic biomass and is, apparently, a congenital property of plants.

Such an innate property can manifest itself in the morphological recapitulation of the properties of the emerging proto-plant in the initial phases of embryogenesis and in the germination of plant seeds in the form of group placing of leucoplasts which became green chloroplasts. For some crops, some mature but partially green seeds germinate poorly, produce weak shoots, and reduce the quality of the product for oilseeds. The reason for this effect can be an excessive and randomly placed chlorophyll, which is not regulated by the genome and, releasing excess oxygen, damages the structures of germinating seeds.

It should be noted that the said above and in (Galitskii 2013, 2016, 2017) was obtained using unique natural data (Tsel’nicker 1994) on the lifetimes of branches of all four orders of spruce (*P. abies* (L.) Karst.) from the Moscow region. Using these four numbers, we solved the following tasks: 1) seven parameters of the spruce dynamics model were determined; 2) the role of initial inhibition of growth and inter-verticillate branches for this species was revealed; 3) a possible cause and mechanism for the appearance of inter-verticillate branches was suggested; 4) model trajectories of the lifespan of branches in the parameter space of sectional model have calculated for spruce and the ruptures of dependence *t^D,j^* (μ) was detected for *j* = 2 and 3 (Fig. 2); 5) the “moments” (the values of μ) of the appearance of branches of all orders in the course of evolution along the parameter μ was determined, including the “moments” for the emergence of the first-order branch in proto-spruce (Figures 3 and 4); 6) the role of group placements of the “points” of photosynthetic biomass in the evolution of the proto-plant was estimated and the corresponding trajectories μ were calculated (Fig. 4). Unfortunately, in the available literature at the moment the data (Tsel’niker 1994) are the only of its kind. It is not difficult to imagine how much progress could been made in researching and understanding the emergence, divergence, and evolution of conifers and other tree species if such data for these species would be available.

In the case of spruce, a possible mechanism for overcoming the rupture of the dependence *t_D,j_* (μ) (Fig. 2) was found using data on inter-verticillate branches (Treskin 1973, Kramer and Kozlovskii 1983, Galitskii 2013). Many other species, in particular conifers, do not have regular inter-verticillate branches, although according to the model they have passed or will have to go through a similar situation of morphological divergence and, accordingly, use some kind of mechanism for bridging the gap. Apparently, when analyzing such situations, a huge array of accumulated results of studies of morphological, physiological, anatomical, biochemical and other features and mechanisms of existing and ancient plants will be in demand and used.

## Acknowledgements

Author thanks, D.O Logofet, V.D. Lahno and V.A. Shuvalov for supporting the work. The research was carried out using supercomputers at Joint Supercomputer Center of the Russian Academy of Sciences (JSCC RAS).

## Footnotes

1 It should be noted that is the *only article* (Tsel’niker 1994) containing two sets of data on the lifetime of branches of all orders of the tree (spruce). These data were used to parameterize the spruce’s model and its further analysis (Galitskii 2013, 2017).

